# RAP1-RHO small GTPase cross-talk mediates integrin-dependent and - independent platelet procoagulant response

**DOI:** 10.1101/2025.05.22.655614

**Authors:** Abigail Ballard-Kordeliski, Nikola Ziegmann, Wyatt Schug, Mark H. Ginsberg, Antje Schaefer, Robert H. Lee, Wolfgang Bergmeier

## Abstract

Platelet adhesion and procoagulant activity are critical for primary and secondary hemostasis, respectively. The small GTPase RAP1 is a central regulator of platelet aggregation as it controls αIIbβ3 integrin activation through direct interaction with the integrin adapter protein, TALIN-1 (Tln-1). In addition to their aggregation defect, activated platelets lacking RAP1 (*Rap1*^*mKO*^) exhibited a marked impairment in surface exposure of phosphatidylserine (PtdSer), a negatively charged phospholipid with procoagulant activity. However, the mechanisms by which RAP1 regulates PtdSer exposure are unclear.

Here we investigated the hypothesis that RAP1 regulates platelet PtdSer exposure through cross-talk with small GTPases of the Rho family. Consistent with their defect in PtdSer exposure, *Rap1*^*mKO*^ platelets showed reduced procoagulant activity *in vitro* and *in vivo* when compared to controls. Stimulated *Rap1*^*mKO*^ platelets exhibited elevated RHOA-GTP levels, and inhibition of the RHOA effector, Rho associated coiled-coil kinase (ROCK), partially restored PtdSer exposure in these cells. A milder defect in PtdSer exposure was observed for platelets from *Tln-1*^*mR35/118E*^ mice, i.e. mice with impaired RAP1-Tln-1 interaction but otherwise intact RAP1 signaling. ROCK inhibition fully restored PtdSer exposure in *Tln-1*^*mR35/118E*^ platelets. Opening of the mitochondrial permeability transition pore, a cellular response critical to PtdSer exposure, was impaired in *Rap1*^*mKO*^ platelets and restored by pretreatment of cells with the ROCK inhibitor.

Our study provides first evidence that platelet RAP1 signaling affects hemostatic plug formation independent of its key role in platelet adhesion. Additionally, our studies strongly suggest that RAP1 regulates PtdSer exposure and procoagulant activity in a RHOA/integrin-dependent and -independent manner.

## Introduction

Within a hemostatic plug there are two classifications of activated platelets: proadhesive and procoagulant.^1^ Pro-adhesive platelets are characterized by high affinity integrin receptors, such as αIIbβ3, which mediate platelet adhesion to the site of injury and platelet aggregation via binding of fibrinogen and other ligands.^2^ Integrin-mediated platelet adhesion is essential to formation of the initial platelet plug to cease bleeding. Procoagulant platelets facilitate thrombin generation and thus enhance the formation of fibrin, a fibrous protein critical for hemostatic plug stability. Characteristic to procoagulant platelets is exposure of the negatively charged phospholipid, phosphatidylserine (PtdSer), on the cell surface.^3^ Procoagulant platelet formation is dependent on sustained high cytosolic calcium levels and mitochondrial depolarization, mediated by the opening of the mitochondrial permeability transition pore (MPTP) via its essential adaptor protein cyclophilin D (CypD).^4,5^ Depolarization of the mitochondria ultimately allows the scramblase transmembrane protein 16 (TMEM16F) to flip PtdSer to the outer plasma membrane leaftlet.^6,7^

Platelet activation is a tightly controlled process with small GTPases playing a central role.^8–10^ Small GTPases are molecular switches which are active in their GTP-bound state and inactive in their GDP-bound state. When GTP bound, small GTPases undergo conformational changes to interact with effector proteins.^11^ The role of small GTPases in αIIbβ3 integrin activation is well-defined, while their activity in procoagulant platelet formation remains less explored.

The Ras family GTPase, RAP1, is the most abundant small GTPase in platelets.^9^ RAP1 is a central regulator of platelet adhesion/aggregation as loss of both isoforms (RAP1a and RAP1b) markedly impairs integrin activation, resulting in significantly prolonged bleeding times.^12^ Activation of αIIbβ3 integrin requires direct interaction between RAP1 and its effector protein Tln-1.^13–15^ Disruption of the interaction between RAP1 and Tln-1 (*Tln-1*^*mR35/118E*^) results in loss of αIIbβ3 integrin activation while retaining other RAP1 signaling responses.^13^ Ligand binding to αIIbβ3 integrin induces outside-in signaling, an important mechanism to potentiate platelet activation.^16,17^ Loss of RAP1 also results in impaired platelet procoagulant response^13,18^, although the mechanism of RAP1 mediated PtdSer exposure is unclear.

The RHO family GTPase, RHOA, is primarily known for its role in cytoskeletal rearrangements.^10,19^ Activation of RHOA occurs downstream of G_13_-coupled receptors, whereas inhibition of RHOA results from integrin outside-in signaling.^20,21^ In contrast to RAP1, inhibition of RHOA signaling leads to increased PtdSer exposure.^22^ Crosstalk between RAP1 and RHOA was demonstrated for other cell types^23,24^; however, the potential role of RAP1-RHOA crosstalk in platelet procoagulant response remains unexplored.

In the present study, we investigated the mechanism of RAP1-mediated PtdSer exposure in platelets. Our studies demonstrate that RAP1-mediated PtdSer exposure occurs through integrin-dependent and -independent mechanisms, and that the integrin-dependent mechanism occurs through a connection to RHOA.

## Results

### Decreased PtdSer exposure and thrombin generation potential in RAP1-deficient platelets in vitro

Dual-agonist stimulation of platelet glycoprotein (GP)VI and protease-activated receptor (PAR) 4 results in robust PtdSer exposure *in vitro*^3^. Therefore, procoagulant response was stimulated with convulxin (CVX; GPVI) and PAR4-activating peptide (PAR4p) and measured via Annexin V binding using flow cytometry (percentage PtdSer-positive events). Compared to controls, exposure of PtdSer was significantly impaired in platelets lacking RAP1 (*Rap1*^*mKO*^) activated with CVX/PAR4p **(Figure 1A)**. Using a modified thrombin generation assay^25^ in which activated procoagulant platelets in platelet-rich plasma (PRP) provide the negatively charged phospholipid surface required for coagulation factor assembly, we evaluated the contribution of RAP1-mediated PtdSer exposure to thrombin generation *in vitro*. Thrombin generation was minimal in platelet poor plasma (PPP) when compared to PRP samples (**Figure 1B,C)**. Congruent with the defect in PtdSer exposure, thrombin generation, defined as peak thrombin (nM), in *Rap1*^*mKO*^ PRP was significantly reduced when compared to control PRP; however, there was no significant difference in time to peak **(Figure 1B-D)**.

**Figure 1.**
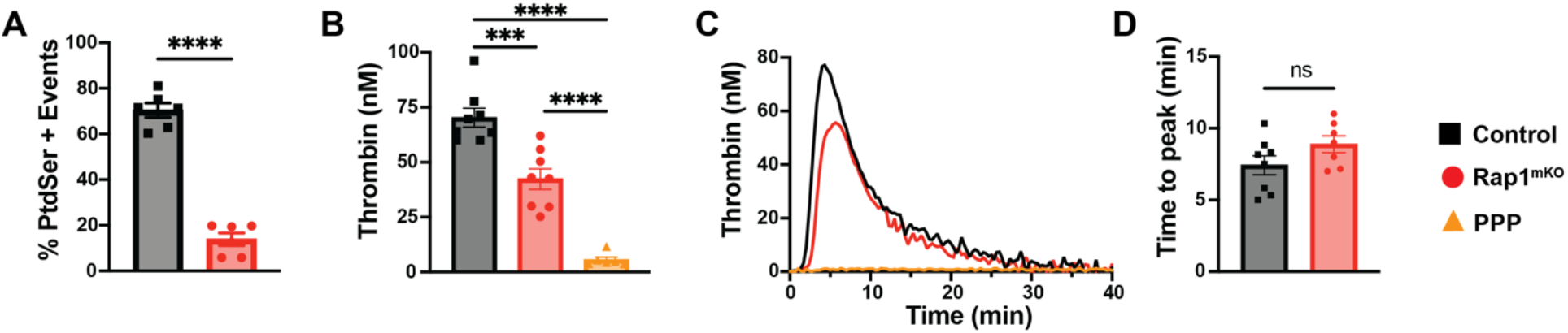
RAP1 mediated PtdSer exposure contributes to thrombin generation *in vitro*. **(A)** Flow cytometry analysis of procoagulant response (% PtdSer+ events (Annexin V binding)) in control and *Rap1*^*mKO*^ platelets stimulated with 50 ng/ml CVX + 250 µM Par4p; n=6. **(B)** Peak thrombin (nM) levels for control platelet rich plasma (PRP), *Rap1*^*mKO*^ PRP, and platelet poor plasma (PPP); n=9. **(C)** Representative thrombin generation curves for indicated samples. **(D)** Time to peak thrombin generation for control and *Rap1*^*mKO*^ PRP; n=7. ***P<0.001, ****P<0.0001, ns: not significant.

### Impaired procoagulant activity of RAP1-deficient platelets in vivo

We recently developed a new imaging model to quantify platelet procoagulant activity during hemostatic plug formation *in vivo*.^26^ In this model, perforating injuries ∼50 μm in diameter are induced to the saphenous vein by laser injury, and the 3-dimensional accumulation of platelets and fibrin is monitored in real time by spinning disk confocal microscopy. Using this model, we were able to demonstrate that fibrin accumulation is significantly impaired in mice lacking CypD in platelets only, demonstrating that platelets are the main procoagulant cellular surface during hemostatic plug formation.^26^ One limitation of the model is that visualizing and measuring hemostatic plug components is difficult in mice with excessive bleeding, including *Rap1*^*mKO*^ mice. To circumvent this limitation and determine the contribution of RAP1 to platelet procoagulant activity *in vivo*, we used an adoptive platelet transfer strategy to generate mice with specific defects in platelet function without excess bleeding: 1) mice that received a mixture of *CypD*^*-/-*^ and WT platelets at a ratio of 3/1, and 2) mice that received a mixture of *CypD*^*-/-*^ and *Rap1*^*mKO*^ platelets at a ratio of 3/1. *CypD*^*-/-*^ platelets were co-transfused to facilitate hemostatic plug formation in the context of RAP1-deficiency **(supplemental Figure 1)**. *CypD*^*-/-*^ platelets, however, lack procoagulant activity in this model.^26^ Thus, fibrin accumulation at sites of injury in co-transfused mice would depend on the procoagulant activity of WT or *Rap1*^*mKO*^ platelets. As shown in **Figure 2**, fibrin accumulation was significantly reduced in mice transfused with *CypD*^*-/-*^*/Rap1*^*mKO*^ platelets when compared to mice transfused with *CypD*^*-/-*^*/*WT platelets. Together, these studies demonstrate a critical role for RAP1 in platelet procoagulant activity *in vitro* and *in vivo*.

**Figure 2.**
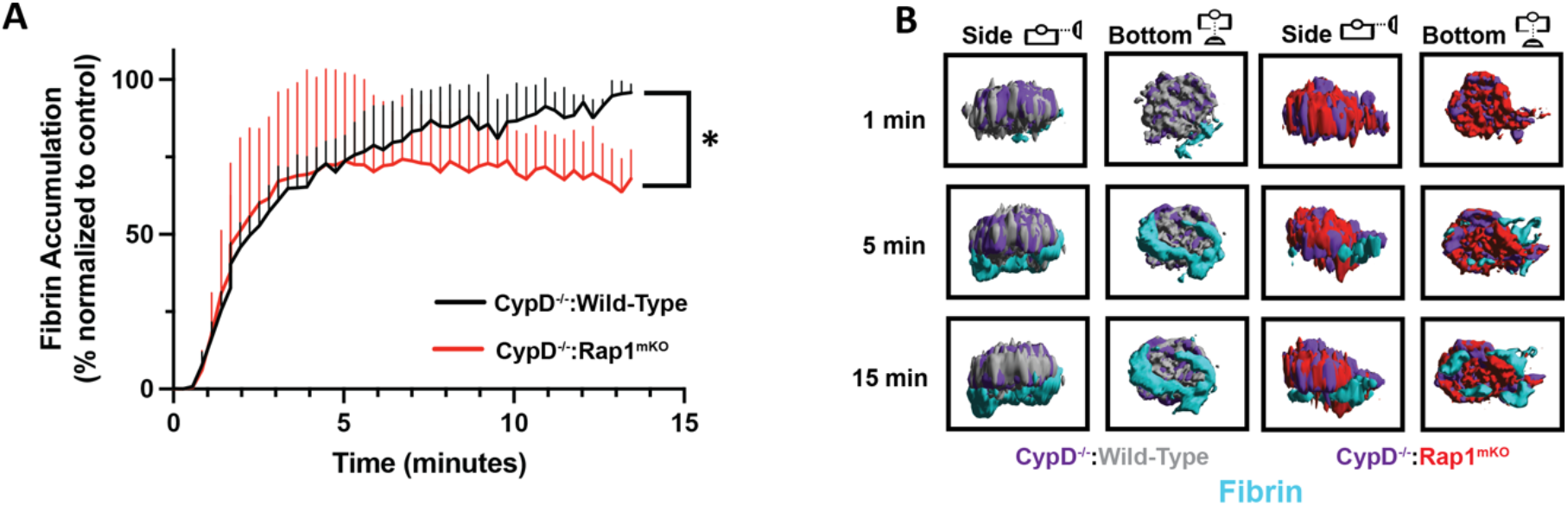
Reduced procoagulant activity of *Rap1*^*mKO*^ platelets *in vivo*. **(A)** Fibrin accumulation at sites of laser injury in mice transfused with *CypD*^*-/-*^ and *Rap1*^*mKO*^ platelets (n=21 injuries; 3 mice) compared to mice transfused with *CypD*^*-/-*^ and wild-type platelets (n=20; 3 mice). Data are shown as sum fluorescence intensity +/-SEM. *P<0.05 **(B)** Representative images of hemostatic plugs in mice transfused with *CypD*^*-/-*^ (purple) and wild-type (grey) or *CypD*^*-/-*^ (purple) and *Rap1*^*mKO*^ (red) platelets. Fibrin is shown in cyan. Images are presented with side and bottom views at indicated time points.

### Integrin-dependent and -independent mechanisms of RAP1-mediated PtdSer exposure

Given the documented role of integrin outside-in signaling in PtdSer exposure^27^, we next determined whether the defect in platelet procoagulant response observed in *Rap1*^*mKO*^ platelets is secondary to their defect in αIIbβ3 integrin activation. We compared the response to dual-agonist stimulation in *Rap1*^*mKO*^ platelets to platelets from mice with impaired Tln-1-RAP1 interaction (*Tln-1*^*mR35/118E*^). Consistent with previous studies with single agonists, both *Rap1*^*mKO*^ and *Tln-1*^*mR35/118E*^ platelets exhibited a marked defect in αIIbβ3 integrin activation (JON/A-PE binding) in response to dual-agonist stimulation **(Figure 3 A,D)**, while granule secretion (P-selectin (α-CD62P) surface expression) was not impaired **(Figure 3 B,E)**. Dual agonist-induced PtdSer exposure was also significantly impaired in *Rap1*^*mKO*^ and *Tln-1*^*mR35/118E*^ platelets **(Figure 3C,F)**. Interestingly, PtdSer exposure was reduced by 85% in *Rap1*^*mKO*^ platelets, while only a 48% reduction in PtdSer-positive cells was observed for *Tln-1*^*mR35/118E*^ platelets compared to controls. Thus, these studies suggested that RAP1 facilitates platelet PtdSer exposure by integrin-dependent and -independent mechanisms.

**Figure 3.**
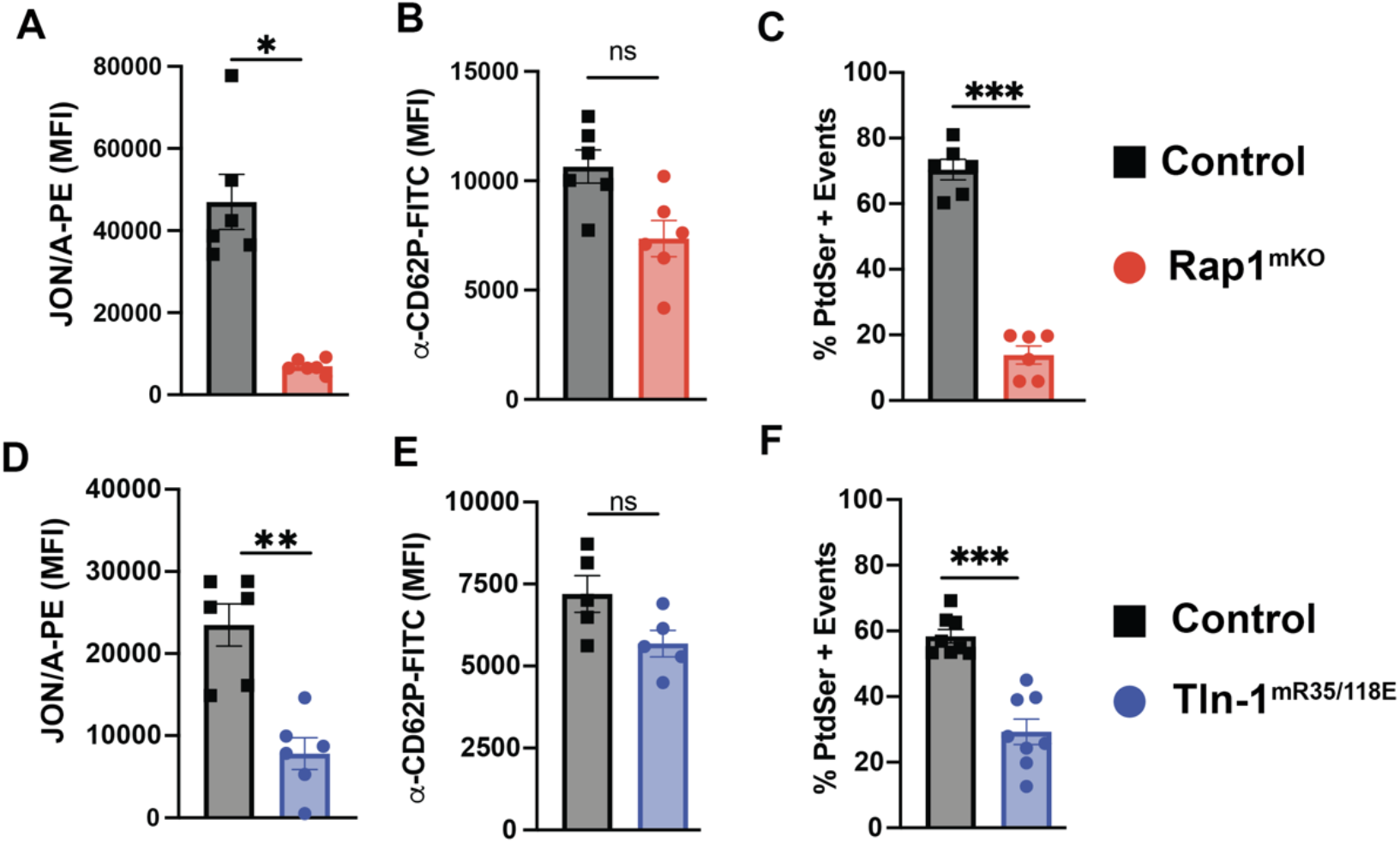
RAP1 affects platelet PtdSer exposure via integrin independent and dependent mechanisms. **(A-C)** Flow cytometry analysis of αIIbβ3 activation (JON/A-PE) (A), granule secretion (α-CD62P-FITC) (B), and PtdSer exposure (Annexin V-AF647) (C) for control or *Rap1*^*mKO*^ platelets stimulated with 50 ng/ml CVX + 250 µM Par4p (n=6). **(D-F)** Flow cytometry analysis of αIIbβ3 activation (JON/A-PE) (D), granule secretion (α-CD62P-FITC) (E), and PtdSer exposure (Annexin V) (F) for control or *Tln-1*^*mR35/118E*^ platelets stimulated with 50 ng/ml CVX + 250 µM Par4p (n=8). *P<0.05, **P<0.01, **P<0.001, ns: not significant.

Integrin outside-in signaling negatively regulates the activation state of the small GTPase RHOA^28^, and previous work demonstrated that inhibition of RHOA signaling leads to increased PtdSer exposure in platelets.^22^ Thus, we hypothesized that impaired PtdSer exposure in *Rap1*^*mKO*^ platelets results, at least in part, from elevated RHOA activity. Consistent with this hypothesis, we observed significantly increased RHOA-GTP levels in *Rap1*^*mKO*^ and *Tln-1*^*mR35/118E*^ platelets at 2 min after addition of agonists **(Figure 4A,B)**. We next studied dual agonist-induced platelet activation in *Rap1*^*mKO*^ and *Tln-1*^*mR35/118E*^ platelets pretreated with an inhibitor of the well-established RHOA effector protein, ROCK.^22^ Treatment with the ROCK inhibitor (YM-27632) did not significantly affect granule secretion or αIIbβ3 activation in controls, *Rap1*^*mKO*^, or *Tln-1*^*mR35/118E*^ platelets **(supplemental Figure 2)**. However, inhibition of ROCK partially recovered PtdSer exposure in *Rap1*^*mKO*^ platelets **(Figure 4C)**. Importantly, ROCK inhibition fully restored PtdSer exposure in *Tln-1*^*mR35/118E*^ platelets **(Figure 4D)**.

**Figure 4.**
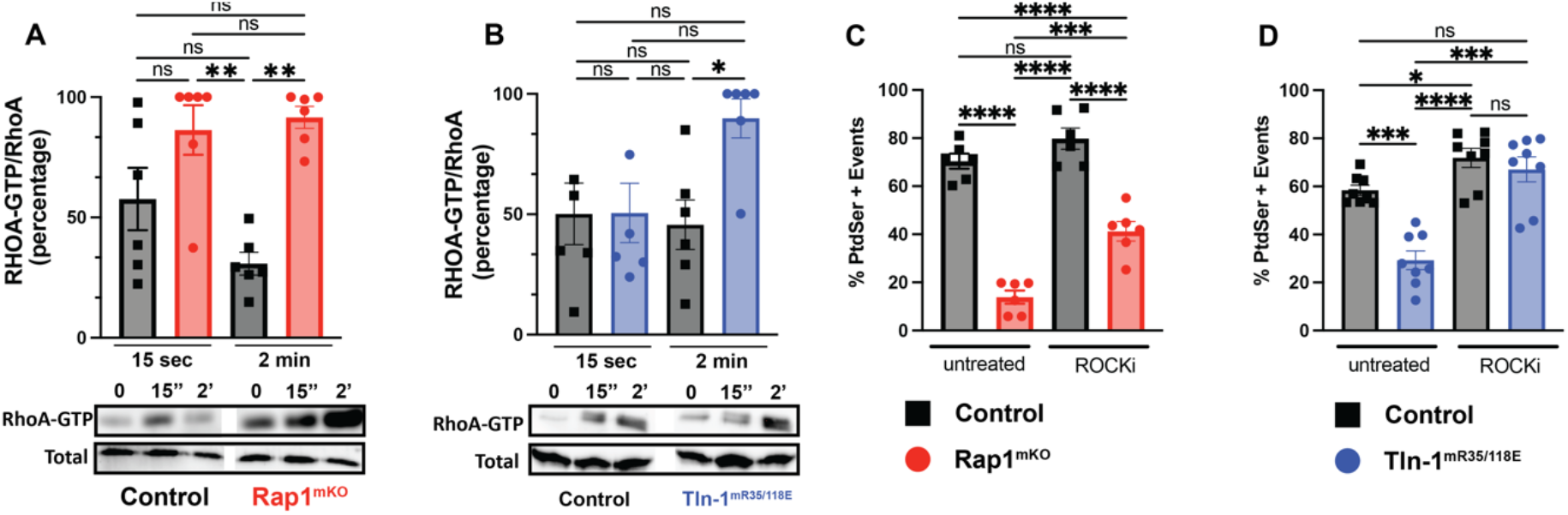
Integrin dependent RAP1-RHOA connection regulates PtdSer exposure. **(A)** Active (GTP-bound) RHOA over total RHOA in control (black bars) or *Rap1*^*mKO*^ platelets (red bars) activated for the indicated times with 50 ng/ml CVX + 250 µM Par4p (n=5). Percentages normalized to highest value. **(B)** Active (GTP-bound) RHOA over total RHOA in control (black bars) or *Tln-1*^*mR35/118E*^ platelets (blue bars) activated for the indicated times with 50 ng/ml CVX + 250 µM Par4p (n=5). Percentages normalized to highest value. **(C)** Flow cytometry analysis of Annexin V binding (% PtdSer + events) on control or *Rap1*^*mKO*^ platelets pretreated with ROCK inhibitor (YM-27632) and then stimulated with 50 ng/ml CVX + 250 µM Par4p (n=6). **(D)** Flow cytometry analysis of Annexin V binding (% PtdSer + events) on control or *Tln-1*^*mR35/118E*^ platelets pretreated with ROCK inhibitor (YM-27632) and then stimulated with 50 ng/ml CVX + 250 µM Par4p (n=8). *<0.05, **<0.01, ***<0.001, ns: not significant

### Calcium mobilization is not affected in Rap1^mKO^ platelets

One of the key factors to successful platelet procoagulant response is a robust and sustained increase in cytosolic calcium levels, which is required to trigger mitochondrial depolarization.^3,5^ To determine cytosolic calcium levels, control or *Rap1*^*mKO*^ platelets were labeled with the calcium-sensitive dye, Fluo-4 ^5^, and activated in the presence or absence of ROCK inhibitor. Dual-agonist stimulation resulted in sustained high calcium levels in control and *Rap1*^*mKO*^ platelets, both in the presence and absence of ROCK inhibitor **(Figure 5A)**. The integrated calcium signal (area under the curve) was comparable between *Rap1*^*mKO*^ and control platelets. Slightly increased calcium levels were observed for both control and *Rap1*^*mKO*^ platelets activated in the presence of ROCK inhibitor **(Figure 5B)**. Under these experimental conditions, *Rap1*^*mKO*^ platelets had decreased PtdSer exposure compared to controls; and inhibition of RHOA/ROCK signaling led to a significant increase in PtdSer-positivity for control and *Rap1*^*mKO*^ platelets, similar to **Figure 4C**. Together, these studies suggest that altered calcium mobilization does not account for the observed differences in PtdSer exposure observed in *Rap1*^*mKO*^ platelets.

**Figure 5.**
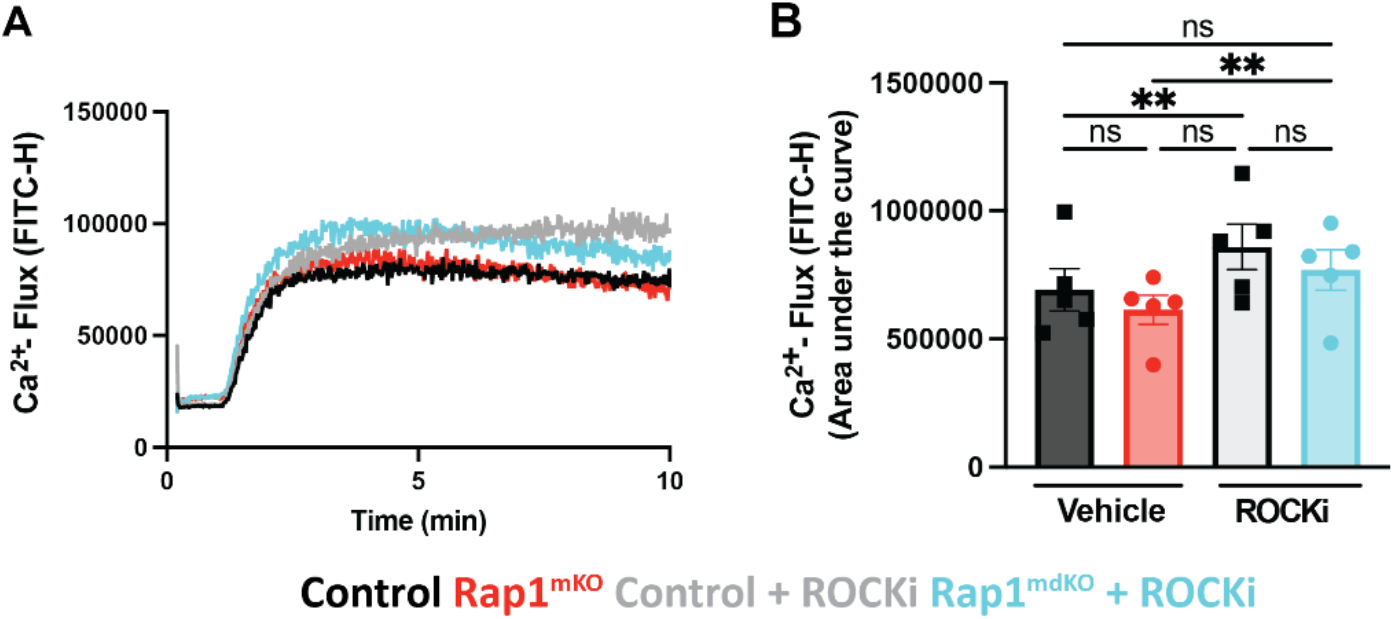
Calcium mobilization is not affected in *Rap1*^*mKO*^ platelets. **(A)** Representative curves for Fluo-4 fluorescence (FITC-H) in control (black and grey curves) or *Rap1*^*mKO*^ platelets (red and blue curves) activated in the presence or absence of ROCK inhibitor (YM-27632) with 50 ng/ml CVX + Par4p 150 µM. **(B)** Area under the curve analysis for Fluo-4 fluorescence traces described in (A) (n=5). *P<0.05, **P<0.01, ***P<0.001, ns, not significant

### Mitochondrial depolarization is decreased in Rap1^mKO^ platelets and partially recovered by inhibition of RHOA/ROCK signaling

Opening of the MPTP and mitochondrial depolarization are critical events for platelet procoagulant response.^4,29^ We next studied mitochondrial depolarization in dual agonist-stimulated platelets using the JC-1 flow cytometry-based assay.^30^ Mitochondrial depolarization was significantly impaired in *Rap1*^*mKO*^ platelets when compared to controls **(Figure 6A,B)**. Pretreatment with ROCK inhibitor led to markedly increased mitochondrial depolarization in both control and *Rap1*^*mKO*^ platelets **(Figure 6A,B)**.

**Figure 6.**
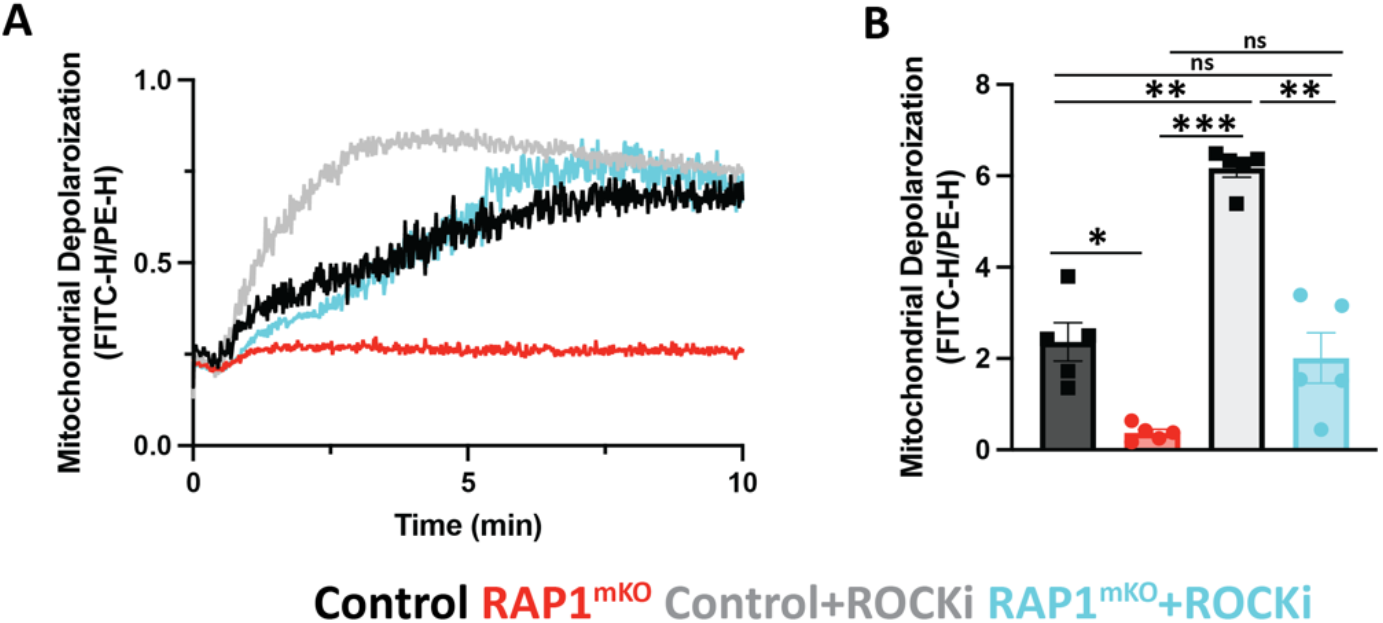
Mitochondrial depolarization is decreased in *Rap1*^*mKO*^ platelets and partially recovered with ROCK inhibitor. **(A)** JC-1 fluorescence ratio (FITC-H/PE-H) for control (black and grey curves) or *Rap1*^*mKO*^ platelets (red and blue curves) activated in the presence or absence of ROCK inhibitor (YM-27632) with 50 ng/ml CVX + Par4p 150 µM. **(B)** Area under the curve (AUC) analysis (n=4). *P<0.05, **P<0.01, ***P<0.001, ns, not significant.

## Discussion

Procoagulant platelets, i.e. platelets exposing PtdSer on their outer plasma membrane, are critical to thrombin/fibrin generation and hemostatic plug stability. Small GTPases of the RAS (RAP1) and RHO (RHOA, RAC1) families are known regulators of PtdSer exposure in platelets. Our study provides first evidence for RAP1-RHO cross-talk required for platelet PtdSer exposure and platelet-dependent thrombin generation, both *in vitro* and *in vivo*.

RAP1 is an essential regulator of αIIbβ3 integrin mediated platelet adhesion and hemostatic plug formation.^12,13^ While no patients with loss-of-function mutations in RAP1 have been identified, mutations in CalDAG-GEFI (*RASGRP2*), an important regulator of RAP1 activation, are known. These mutations cause moderate to severe bleeding in humans^31–33^, similar to what was shown for mice deficient in RAP1 or CalDAG-GEFI.^12,34^ As outlined above, RAP1 signaling is also important for procoagulant platelet formation. Importantly, bleeding is also observed in patients with Scott Syndrome, i.e. in patients with a defect in procoagulant platelet formation but not αIIbβ3 integrin activation.^35^ Scott Syndrome is attributed to mutations in TMEM16F, a phospholipid scramblase that mediates PtdSer translocation to the outer membrane layer.^7,36^ TMEM16F is highly expressed in platelets but is also found in other cell types including erythrocytes and endothelial cells (ECs).^37–39^ The similar bleeding phenotype between TMEM16F germline and megakaryocyte/platelet-specific knockout mice suggests that impaired procoagulant activity in platelets is the main reason for bleeding observed in Scott syndrome patients.^6,7,37^ Furthermore, we recently demonstrated a key role of procoagulant platelets, but not ECs for thrombin/fibrin formation during hemostatic plug formation.^26^ Here we provide the first evidence that RAP1 mediated platelet PtdSer exposure contributes to thrombin/fibrin generation *in vitro* and *in vivo*. These studies suggest that bleeding in patients with impaired RAP1 signaling may result in part from reduced thrombin/fibrin formation at sites of vascular injury.

The RHO family of GTPases plays an important role in platelet pro-coagulant response. RHOA/ROCK signaling inhibits PtdSer exposure^22,27,40^ and thus needs to be downregulated by integrin outside-in signaling during platelet pro-coagulant response.^27,28^ We observed prolonged RHOA activation in *Rap1*^*mKO*^ and *Tln-1*^*mR35/118E*^ platelets, i.e. platelets with markedly impaired integrin inside-out activation. Our studies further established that inhibition of RHOA/ROCK signaling fully restored PtdSer exposure in *Tln-1*^*mR35/118E*^ platelets, which are defective in RAP1-TALIN interaction. However, only a partial recovery of PtdSer exposure by inhibition of RHOA/ROCK signaling was observed in *Rap1*^*mKO*^ platelets, a finding that suggests a TALIN1/integrin/RHOA-independent contribution of RAP1 to platelet procoagulant response. We previously showed that in platelets RAP1 positively regulates the activity of RAC1^12,41^, another RHO GTPase with a documented role in PtdSer exposure^22^. Importantly, RAC1 and RHOA are also known to negatively regulate each other’s activation state.^42^ Together, these studies suggest an important role for cross-talk between RAP1 and RHO GTPases in integrin-dependent and -independent PtdSer exposure.

High sustained cytosolic calcium levels and mitochondrial depolarization are two critical steps in procoagulant platelet formation. Inhibition of RHOA/ROCK signaling led to a small but significant increase in cytosolic calcium and a marked increase in mitochondrial depolarization in both control and *Rap1*^*mKO*^ dual agonist-stimulated platelets. Compared to controls, a significant decrease in mitochondrial depolarization but not calcium mobilization was observed for dual agonist-stimulated RAP1-deficient platelets. Given that calcium mobilization was not altered in *Rap1*^*mKO*^ platelets, another mechanism must account for the decreased mitochondrial depolarization. Reactive oxygen species also affect mitochondrial depolarization, and both RHOA and RAC1 have been shown to regulate ROS production.^43^ Whether impaired mitochondrial depolarization and PtdSer exposure in *Rap1*^*mKO*^ platelets is caused by dysregulated ROS production will be a topic of future studies.

In conclusion, our studies demonstrate that impaired RAP1 signaling leads to decreased platelet procoagulant response and thrombin generation *in vitro* and *in vivo*. RAP1 affects PtdSer exposure via integrin-dependent and -independent mechanisms, which likely involve cross-talk with RHO GTPases and the depolarization of the mitochondrial permeability transition pore. This study improves our understanding of the role of small GTPases in platelet procoagulant response and thus may have important implications for the development of better therapies to prevent bleeding or thrombosis.

## Acknowledgements

The authors thank David Paul and Summer Jones (University of North Carolina at Chapel Hill) for valuable technical support and valuable expertise. This work was supported by the National Institutes of Health, National Heart, Lung, and Blood Institute (grants R35 HL144976 [W.B.], F31 HL165935 [A.B.-K.], P01 HL151433 [M.H.G. and W.B.])

## Author Contributions

A.B.-K. designed research, performed research, analyzed data and wrote manuscript. N.Z. and W.S. performed research and analyzed data. A.J designed and performed research. R.H.L designed research. M.H.G. contributed vital regents. W.B designed research and wrote manuscript.

## Methods

### Mice

Generation of *Rap1*^*mKO* 12^, *Tln-1*^*mR35/118E* 13^, and IL4Rα-GPIbα-tg ^44^ mice has been previously described. Experimental procedures were approved by the Institutional Animal Care and Uses Committee.

### 4D Saphenous Vein Laser Injury Model

Adoptive transfer of platelets into thrombocytopenic mice was performed as previously described.^26,45,46^ In brief, blood was collected in PBS with heparin and platelets were washed. Platelets were depleted with α-hIL4R antibody (2.5 μg/g body weight) in IL4Rα/GPIbα-tg mice. Washed platelets were labeled with Alexa488 or Alexa647-labeled antibodies to GPIX and administered via retroorbital injection into platelet depleted IL4Rα/GPIbα-tg mice to a final circulating concentration of 2-5 × 10^8^ platelets/ml. 4D imaging of saphenous vein laser injury was performed as recently described.^26^ In brief, the saphenous vein was exposed and relevant fluorescently labeled antibodies were administered via retroorbital injection. Laser ablation was used to create a perforating injury in the vein and spinning disk confocal imaging was performed on a Zeiss Axio Examiner Z1 inverted spinning disk confocal microscope (4 × 4 binning, 7.5 um step, 150 um total travel). Images were acquired with SLIDEBOOK 6.0 software (Intelligent Imaging Innovations). Image analysis was performed using ImageTank software (Visual Data Tools, Inc) as previously described.^22^

### Flow Cytometry

Platelets were washed as previously described^12^ and activated with the indicated concentrations of convulxin ((CVX) (purchased from Kenneth Clemetson, Theodor Kocher Institute, University of Berne, Switzerland) and Par4p (GL Biochem) in the presence of 2 μg/ml JON/A-PE (clone Leo.H4, Emfret Analytics), α-P-selectin-FITC (clone RB40.34, BD Biosciences), and Annexin V-AF647. Where indicated, samples were incubated with ROCK inhibitor (YM-27632) for 10 minutes prior to activation. After 15 minutes of incubation, samples were diluted and analyzed via flow cytometry (Accuri C6 Plus flow cytometer (BD Biosciences)).

### RAP1 and RHOA activation assays

Washed platelets (260 μL samples at 8 × 10^8^/ml) were stimulated with 50 μg/ml CVX and 250 μm Par4p in aggregometry. Platelets were lysed with cold 2x lysis buffer (100 mmol/L Tris/HCl pH 7.4, 400 mmol/L NaCl, 5 mmol/L MgCl2, 2% Nonidet P-40, 20% glycerol and protease inhibitor cocktail lacking ethylenediaminetetraacetic acid (Roche)). Lysates were incubated with RalGDS-RBD beads (Millipore, Billerica, MA) for RAP1-GTP or Rhotekin-RBD beads (Cytoskeleton) for RHOA-GTP for 1 hour at 4°C. Beads were washed 3 times then resuspended in 2x Laemmli buffer for detection of RAP1-GTP or RHOA-GTP via western blotting via standard western blotting procedure. For loading controls, 50 μL of the platelet sample was combined with 50 μL 2x Laemmli buffer (75 mmol/L Tris/HCl, pH 6.8, 2% sodium dodecyl sulfate, 10% glycerol, 5% 2-mercaptoethanol, 0.002% bromophenol blue). Antibodies to RAP1 (clone 121, Santa Cruz, Cat # sc-65) or RHOA (clone 55, Sigma, Cat # 05-778) were used for detection of RAP1 or RHOA, respectively.

### Thrombin generation assay

Thrombin generation assay was performed in a 96-well plate. 0.5 pM TF (Innovin) and 200 ng/ml CVX were added, and wells were brought to volume (30 µl) with Tyrodes buffer containing 1 mM CaCl_2_. Platelet poor plasma (PPP) or platelet rich plasma (PRP, platelet count 5 × 10^8^/ml) was added to initiate the reaction. Each sample was calibrated with α2-macroglobulin-thrombin complex calibrator ((Diagnostica Stago Inc Fluca Kit). Fluorescence was measured on a Fluoroskan Ascent fluorometer (Thermo Fisher Scientific, Waltham, MA) with the Ascent Software (version 2.6, Thermo Fisher Scientific) at 37°C.

### JC-1 Mitochondrial Depolarization Assay

Platelets were washed as previously described ^12^ and adjusted to 7.5 × 10^8^/ml. JC-1 dye (Invitrogen, Cat # (1 μl) and washed platelets (20 μl) were added to 180 μl of Tyrode’s buffer with BSA and incubated at 37°C for 10 minutes. 30 μl of labeled platelets were transferred to 70 μl of Tyrode’s buffer containing 2 mM CaCl_2_. Platelets were activated with 100 μl of 2x agonist and 2.5 μg/ml Annexin V-AF647 in Tyrode’s buffer containing 2 mM CaCl_2_, and fluorescence intensities were recorded on a BD Accuri C6 Flow Cytometer. Analysis was completed using FlowJo (FlowJo LLC).

### Fluo-4 Calcium Mobilization Assay

Platelets were washed as previously described ^12^ and adjusted to 1 × 10^9^/ml. Fluo-4 dye was diluted to a concentration of 0.5 mM in DMSO. Platelets (1 × 10^8^/ml in 200 μl Tyrode’s buffer) were labeled with 1 μl of Fluo-4 dye for 30 min at 37°C. After incubation, 800 μl of Tyrode’s buffer was added to labeling reaction. Labeled platelets were diluted (1:1) in Tyrode’s buffer with 4 mM CaCl_2_. Cellular stimulation was induced by addition of a 2x agonist solution. Fluorescence intensity (Fl1) was recorded on a BD Accuri C6 Flow Cytometer. Analysis was completed using FlowJo (FlowJo LLC).

### Statistics

Results are shown as mean +/-standard error of the mean (SEM). Unless otherwise indicated, statistical significance was analyzed via Welch’s t-test.

## References

1. Welsh, J. D. et al. Hierarchical organization of the hemostatic response to penetrating injuries in the mouse macrovasculature. Journal of Thrombosis and Haemostasis 15, 526–537 (2017).

2. Kulkarni, S. et al. A revised model of platelet aggregation. J. Clin. Invest. 105, 783– 791 (2000).

3. Reddy, E. C. & Rand, M. L. Procoagulant Phosphatidylserine-Exposing Platelets in vitro and in vivo. Front. Cardiovasc. Med. 7, 15 (2020).

4. Jobe, S. M. et al. Critical role for the mitochondrial permeability transition pore and cyclophilin D in platelet activation and thrombosis. Blood 111, 1257–1265 (2008).

5. Abbasian, N., Millington-Burgess, S. L., Chabra, S.Malcor, J.-D. & Harper, M. T. Supramaximal calcium signaling triggers procoagulant platelet formation. Blood Advances 4, 154–164 (2020).

6. Fujii, T., Sakata, A., Nishimura, S., Eto, K. & Nagata, S. TMEM16F is required for phosphatidylserine exposure and microparticle release in activated mouse platelets. Proc. Natl. Acad. Sci. U.S.A. 112, 12800–12805 (2015).

7. Suzuki, J., Umeda, M., Sims, P. J. & Nagata, S. Calcium-dependent phospholipid scrambling by TMEM16F. Nature 468, 834–838 (2010).

8. Stefanini, L. & Bergmeier, W. Small GTPases in megakaryocyte and platelet biology. Platelets 30, 7–8 (2019).

9. Stefanini, L. & Bergmeier, W. RAP1-GTPase signaling and platelet function. J Mol Med 94, 13–19 (2016).

10. Aslan, J. E. & Mccarty, O. J. T. Rho GTPases in platelet function. Journal of Thrombosis and Haemostasis 11, 35–46 (2013).

11. Cherfils, J. & Zeghouf, M. Regulation of Small GTPases by GEFs, GAPs, and GDIs. Physiological Reviews 93, 269–309 (2013).

12. Stefanini, L. et al. Functional redundancy between RAP1 isoforms in murine platelet production and function. Blood 132, 1951–1962 (2018).

13. Lagarrigue, F. et al. Talin-1 is the principal platelet Rap1 effector of integrin activation. Blood 136, 1180–1190 (2020).

14. Nieswandt, B. et al. Loss of talin1 in platelets abrogates integrin activation, platelet aggregation, and thrombus formation in vitro and in vivo. The Journal of Experimental Medicine 204, 3113–3118 (2007).

15. Petrich, B. G. et al. Talin is required for integrin-mediated platelet function in hemostasis and thrombosis. The Journal of Experimental Medicine 204, 3103–3111 (2007).

16. Gong, H. et al. G Protein Subunit Gα 13 Binds to Integrin α IIb β 3 and Mediates Integrin “Outside-In” Signaling. Science 327, 340–343 (2010).

17. Shattil, S. J. & Newman, P. J. Integrins: dynamic scaffolds for adhesion and signaling in platelets. Blood 104, 1606–1615 (2004).

18. Lee, R. H., Rocco, D. J., Nieswandt, B. & Bergmeier, W. The CalDAG-GEFI/Rap1/αIIbβ3 axis minimally contributes to accelerated platelet clearance in mice with constitutive store-operated calcium entry. Platelets 34, 2157383 (2023).

19. Schaefer, A. & Der Channing. RHOA takes the RHOad less traveled to cancer. Trends Cancer 8, 655–669 (2022).

20. Pleines, I. et al. Megakaryocyte-specific RhoA deficiency causes macrothrombocytopenia and defective platelet activation in hemostasis and thrombosis. Blood 119, 1054–1063 (2012).

21. Vogt, S., Grosse, R., Schultz, G. & Offermanns, S. Receptor-dependent RhoA Activation in G12/G13-deficient Cells. Journal of Biological Chemistry 278, 28743– 28749 (2003).

22. Dasgupta, S. K. et al. Rho Associated Coiled-Coil Kinase-1 Regulates Collagen-Induced Phosphatidylserine Exposure in Platelets. PLoS ONE 8, e84649 (2013).

23. Kim, J.-G. et al. Ras-related GTPases Rap1 and RhoA Collectively Induce the Phagocytosis of Serum-opsonized Zymosan Particles in Macrophages. Journal of Biological Chemistry 287, 5145–5155 (2012).

24. Cullere, X. et al. Regulation of vascular endothelial barrier function by Epac, a cAMP-activated exchange factor for Rap GTPase. Blood 105, 1950–1955 (2005).

25. Depasse, F. et al. Thrombin generation assays are versatile tools in blood coagulation analysis: A review of technical features, and applications from research to laboratory routine. Journal of Thrombosis and Haemostasis 19, 2907–2917 (2021).

26. Ballard-Kordeliski, A. et al. 4D intravital imaging studies identify platelets as the predominant cellular procoagulant surface in a mouse hemostasis model. Blood 144, 1116–1126 (2024).

27. Pang, A. et al. Shear-induced integrin signaling in platelet phosphatidylserine exposure, microvesicle release, and coagulation. Blood 132, 533–543 (2018).

28. Zhang, Y. et al. Integrin β3 directly inhibits the Gα13-p115RhoGEF interaction to regulate G protein signaling and platelet exocytosis. Nat Commun 14, 4966 (2023).

29. Basso, E. et al. Properties of the Permeability Transition Pore in Mitochondria Devoid of Cyclophilin D. Journal of Biological Chemistry 280, 18558–18561 (2005).

30. Sivandzade, F., Bhalerao, A. & Cucullo, L. Analysis of the Mitochondrial Membrane Potential Using the Cationic JC-1 Dye as a Sensitive Fluorescent Probe. BIO-PROTOCOL 9, (2019).

31. Canault, M. et al. Human CalDAG-GEFI gene (RASGRP2) mutation affects platelet function and causes severe bleeding. Journal of Experimental Medicine 211, 1349– 1362 (2014).

32. Kato, H. et al. Human CalDAG-GEFI deficiency increases bleeding and delays αIIbβ3 activation. Blood 128, 2729–2733 (2016).

33. Lozano, M. L. et al. Novel mutations in RASGRP2, which encodes CalDAG-GEFI, abrogate Rap1 activation, causing platelet dysfunction. Blood 128, 1282–1289 (2016).

34. Crittenden, J. R. et al. CalDAG-GEFI integrates signaling for platelet aggregation and thrombus formation. Nat Med 10, 982–986 (2004).

35. Satta, N., Toti, F., Fressinaud, E., Meyer, D. & Freyssinet, J. M. Scott syndrome: an inherited defect of the procoagulant activity of platelets. Platelets 8, 117–124 (1997).

36. Yang, H. et al. TMEM16F Forms a Ca2+-Activated Cation Channel Required for Lipid Scrambling in Platelets during Blood Coagulation. Cell 151, 111–122 (2012).

37. Baig, A. A. et al. TMEM16F-Mediated Platelet Membrane Phospholipid Scrambling Is Critical for Hemostasis and Thrombosis but not Thromboinflammation in Mice— Brief Report. ATVB 36, 2152–2157 (2016).

38. Liang, P. et al. Deciphering and disrupting PIEZO1-TMEM16F interplay in hereditary xerocytosis. Blood 143, 357–369 (2024).

39. Schmaier, A. A. et al. TMEM16E regulates endothelial cell procoagulant activity and thrombosis. Journal of Clinical Investigation 133, e163808 (2023).

40. Aslan, J. E. et al. The PAK system links Rho GTPase signaling to thrombin-mediated platelet activation. American Journal of Physiology-Cell Physiology 305, C519–C528 (2013).

41. Stefanini, L. et al. Rap1-Rac1 Circuits Potentiate Platelet Activation. ATVB 32, 434– 441 (2012).

42. Nguyen, L. K., Kholodenko, B. N. & Von Kriegsheim, A. Rac1 and RhoA: Networks, loops and bistability. Small GTPases 9, 316–321 (2018).

43. Akbar, H., Duan, X., Saleem, S., Davis, A. K. & Zheng, Y. RhoA and Rac1 GTPases Differentially Regulate Agonist-Receptor Mediated Reactive Oxygen Species Generation in Platelets. PLoS ONE 11, e0163227 (2016).

44. Kanaji, T., Russell, S. & Ware, J. Amelioration of the macrothrombocytopenia associated with the murine Bernard-Soulier syndrome. Blood 100, 2102–2107 (2002).

45. Bergmeier, W. & Boulaftali, Y. Adoptive transfer method to study platelet function in mouse models of disease. Thrombosis Research 133, S3–S5 (2014).

46. Getz, T. M. et al. Novel mouse hemostasis model for real-time determination of bleeding time and hemostatic plug composition. Journal of Thrombosis and Haemostasis 13, 417–425 (2015).

