## supplemental figures for "RAP1-RHO small GTPase cross-talk mediates integrin-dependent and - independent platelet procoagulant response"

### Supplemental Material

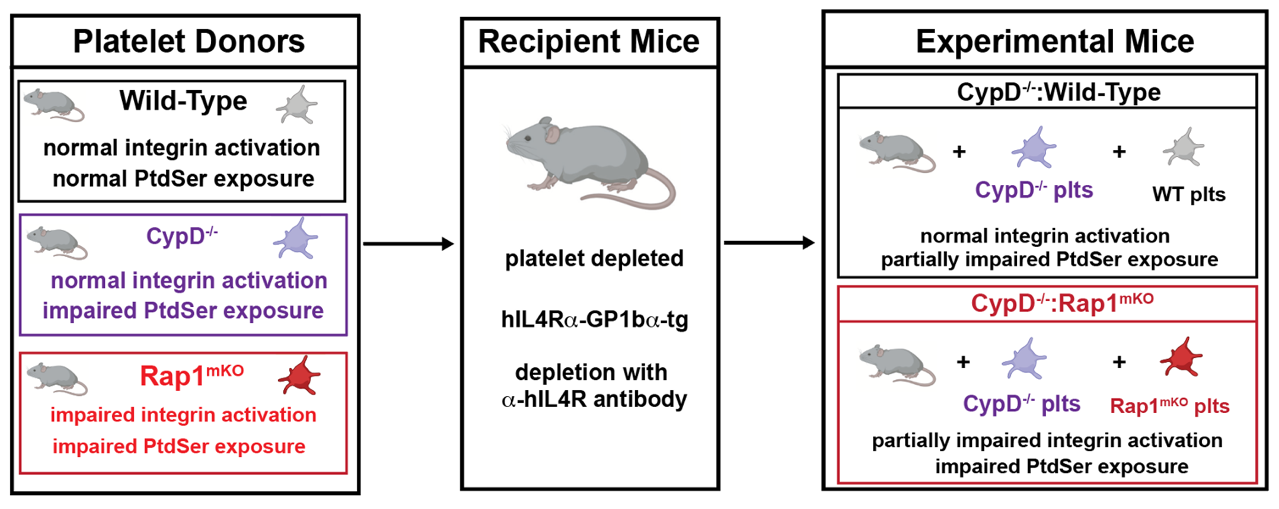

**Supplemental figure 1. Schematic representation for adoptive transfer experiment*.***

Modified platelet adoptive transfer approach.^45^ Platelets from *CypD^-/-^* and wild-type or *CypD^-/-^* and *Rap1^mKO^* mice were transfused into a thrombocytopenic recipient mouse.

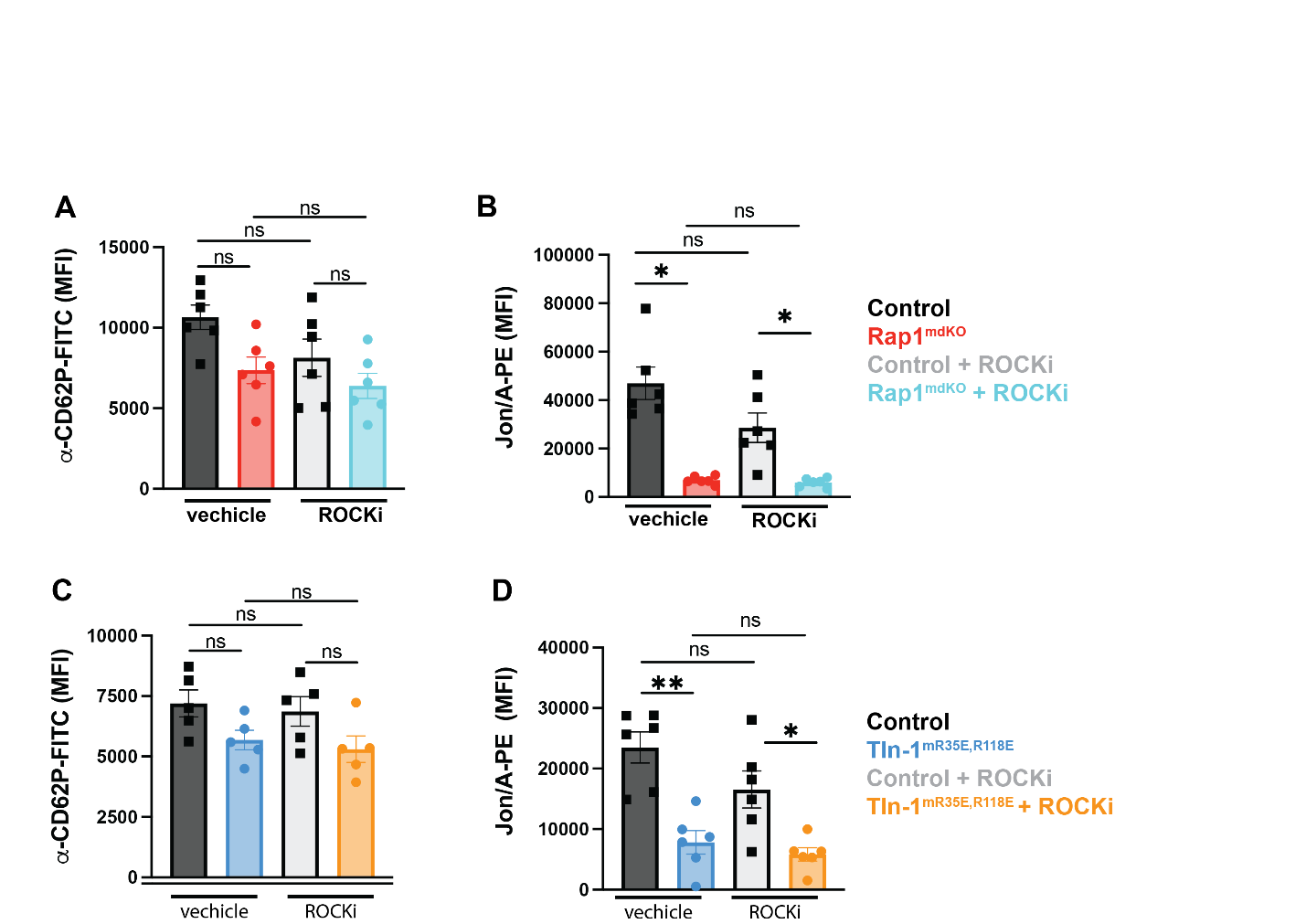

**Supplemental figure 2. ROCK inhibition does not affect granule secretion or αIIbβ3 activation.**

**(A)** Flow cytometry analysis of granule secretion (α-CD62P-FITC) in control (black/grey bars) or *Rap1^mKO^* platelets (red/cyan bars) stimulated with 50 ng/ml CVX + 250 µM Par4p in the presence or absence of ROCK inhibitor (n=6). **(B)** Flow cytometry analysis of αIIbβ3 integrin activation (JON/A-PE) in control or *Rap1^mKO^* platelets stimulated with 50 ng/ml CVX + 250 µM Par4p in the presence or absence of ROCK inhibitor (n=6). **(C)** Flow cytometry analysis of granule secretion (a-CD62P-FITC) in control (black/grey bars) or *Tln-1^mR35/118E^* platelets (blue/yellow bars) stimulated with 50 ng/ml CVX + 250 µM Par4p in the presence or absence of ROCK inhibitor (n=6). **(D)** Flow cytometry analysis of αIIbβ3 integrin activation (JON/A-PE) in control or *Tln-1^mR35/118E^* platelets stimulated with 50 ng/ml CVX + 250 µM Par4p in the presence or absence of ROCK inhibitor (n=6).
